# Cellular basis for cortical network aging in primates

**DOI:** 10.1101/2025.07.08.663725

**Authors:** Melina Tsotras, Joey A. Charbonneau, Claude Lepage, Jeffrey L. Bennett, Jelle Veraart, Alan C. Evans, Eliza Bliss-Moreau, Erika P. Raven

## Abstract

Large-scale brain networks are vulnerable to change with aging and become dysregulated. How these networks are altered at the cellular level remains unclear owing to challenges of bridging data across scales. Here, we integrate *in vivo* cortical similarity networks with whole brain spatial transcriptomics to characterize the aging brain in a lifespan cohort of macaques (N=64, ages 1–26 years). Deep-layer excitatory neurons and oligodendrocytes emerged as dominant correlates of cortical similarity, linking infragranular cell type composition to macroscopic network structure. Age-related declines in network strength were most pronounced in transmodal networks, including default mode and limbic, and aligned with regions enriched in inhibitory and glial cell types. Parvalbumin-enriched chandelier cells showed the strongest association with regional vulnerability, suggesting a role in network disconnection. Cell-type enrichment was conserved across species, with both human and macaque transcriptomic data aligning with the cortical functional hierarchy. These findings uncover a cellular basis for cortical network aging and highlight the value of imaging-transcriptomic integration across scales.

## 1. Introduction

The primate brain contains a diverse array of neuronal and glial cell types, spatially distributed across distinct regions, and embedded within coordinated structural and functional networks ^1–3^. This multiscale architecture scaffolds large-scale brain systems that shape information processing and behavior ^4,5^. Recent studies have uncovered convergence across gene expression, cell types, and neurotransmitter distributions which align with a shared sensory-association hierarchy ^3,6–11^ . While this convergence has been documented, we still lack a mechanistic understanding of how these multiscale features interact, change over time, or predict brain health ^2,12^. Aging is a strong driver of neurobiological change ^13–19^, but the cellular mechanisms underlying age-related reorganization of the cortex remain poorly defined. This is due, in large part, to the challenges of connecting data across spatial and methodological scales. Molecular, cellular, and network-level features often shift together and are interrelated, highlighting the need for integrative approaches to bridge scales and enable in vivo investigation of brain aging.

Traditional methods for studying brain structure in living animals and humans typically use fiber tractography from diffusion MRI ^20–22^ or on morphological features such as cortical thickness. These approaches have limitations, as diffusion tractography can produce false or incomplete maps of connections ^23^ and morphological brain features often lack sensitivity to more subtle microstructural changes that precede morphological changes. Recent neuroimaging work has shifted towards multivariate, MRI-based analyses that examine how similar brain regions are based on an integration of structural and microstructural features. In this study, we used one such approach called the Morphometric Inverse Divergence (MIND) ^24^. MIND offers a quantitative method for investigating cortical architecture and structural connectivity based on shared micro- and macrostructural properties ^24–26^. Regions with similar structural profiles (e.g, cortical thickness, curvature, myelination) tend to have stronger anatomical connectivity, allowing for MIND-based estimation of cortical similarity networks ^27–29^. MIND has been used across species, including humans, macaques, and rats, and biologically validated using human post mortem data ^24,26,30,31^. In macaques, MIND’s strong association with tract-tracing studies^24^ supports its validity for reflecting true anatomical connectivity. In humans, MIND-based maps align with gene expression patterns^24,30^. These cross-species findings highlight MIND as a powerful tool for linking brain network architecture and cell composition across spatial scales.

Despite advances in multiparametric MRI methods, a gap remains between large-scale brain networks and the underlying cellular architecture of the cortex. This gap limits our ability to resolve how cell types vary across cortical layers and regions. Advances in spatial transcriptomics now allow high-resolution mapping of cell types by assigning molecular signatures directly to anatomy. Studies in humans and nonhuman primates have shown that cortical organization follows consistent developmental and functional gradients, suggesting that spatial rules shape where specific cell types are located^4,32–35^. Most recently, computational advances have allowed the development of whole-brain Stereo-seq atlases, first in mouse^36,37^ and now in cynomolgus macaque^38^. These datasets offer unprecedented opportunities to systematically examine how glutamatergic, GABAergic, and non-neuronal cell types are distributed, how these distributions relate to brain networks, and ultimately examine the cellular basis of age-related cortical reorganization ^38^.

Understanding how cell types are spatially organized across functional brain networks provides key insight into the biological architecture that underlies cortical specialization. This is especially critical in higher-order transmodal cortical regions and networks, where age-related decline is well documented, but its biological basis remains unclear. Here, we link in vivo MRI-derived network organization to ex vivo spatial transcriptomic maps in the primate brain. Using a lifespan cohort of rhesus macaques (N=64, ages 1–26 years) and a recently released whole-brain Stereo-seq atlas ^38^, we identify network-specific cellular profiles and show that regions enriched for non-neuronal and inhibitory neurons exhibit the greatest structural vulnerability with age. We also show the macaque cortex shares fundamental organizational principles with the human brain, enabling cross-species comparison of cell type distributions using a common parcellation framework. These findings provide a multiscale, cross-species model of cortical aging, and demonstrate how cellular composition shapes large-scale network architecture and its susceptibility to change across the lifespan.

## 2. Results

### 2.1 Deep layer excitatory neurons and oligodendrocytes are associated with MRI-derived cortical similarity

The need to understand how MR imaging parameters of the brain relate to actual histological features such as cell type and organization has been clear since the advent of MR imaging ^39–41^. Subject-specific in vivo MRI data of 64 rhesus macaques, cortical atlases, and transcriptomic data (Stereo-seq) were transformed into a common template space ^42,43^ using nonlinear-registration transformations from the CIVET-macaque pipeline. Morphometric INverse Divergence (MIND) was computed *for each subject* to estimate cortical similarity between pairs of cortical regions using seven MRI-derived features (T1w/T2w ratio, cortical thickness, sulcal depth, surface area, gray matter volume, mean curvature, gaussian curvature). Multivariate similarity of these features was computed using Kullback-Leibler Divergence. We computed the *total similarity strength* by averaging the cortical similarity across all regions. We then evaluated the uni- and multivariate spatial correlations between transcriptomic data and total similarity strengths on a cohort-level (see Fig 1, Methods Sections 4.9 & 4.10). Spin permutation tests were performed to control for spatial autocorrelation across cortical regions.

**Figure 1.**
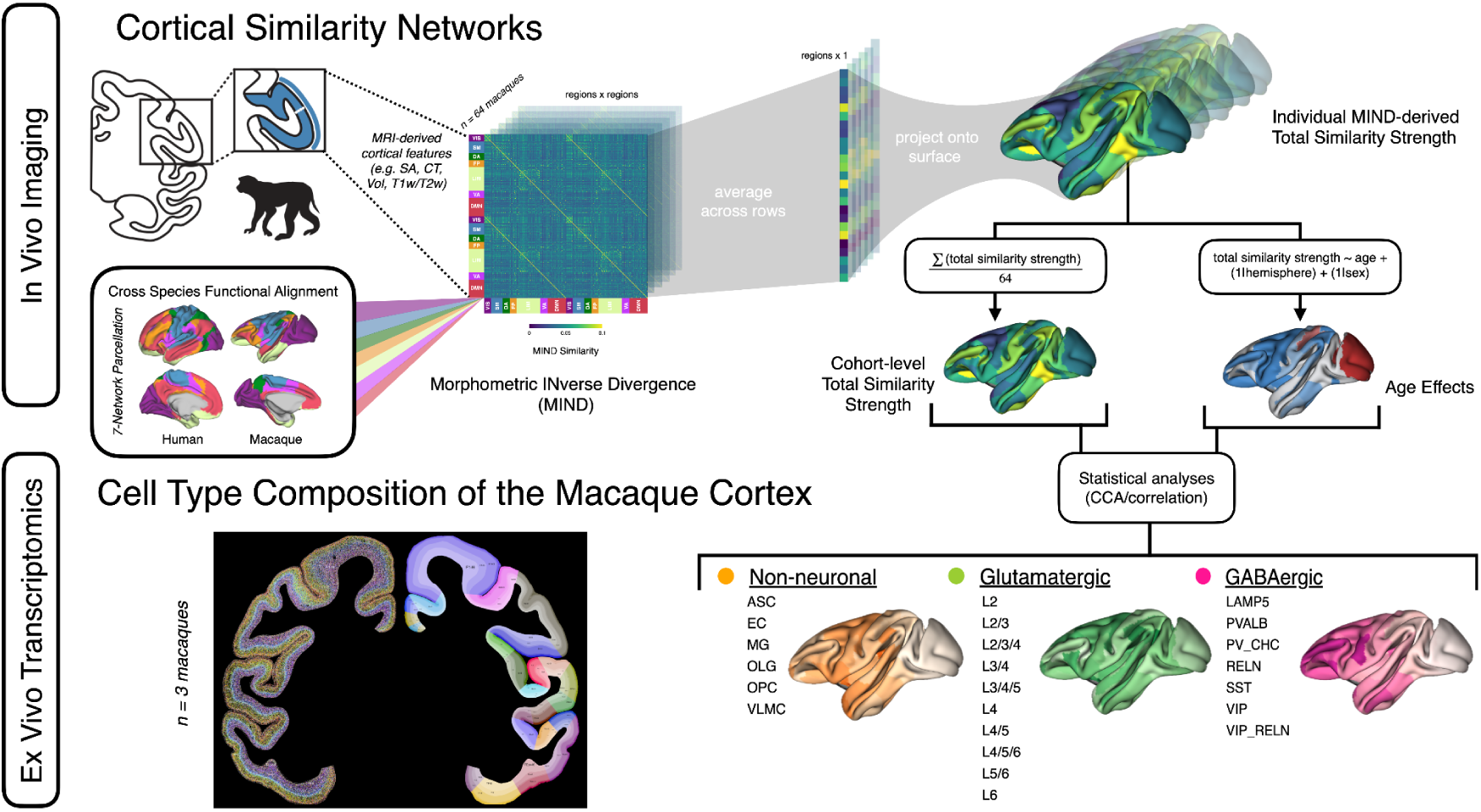
Workflow. In Vivo Imaging. Morphometric INverse Divergence (MIND) was used to estimate subject-level cortical similarity networks using seven MRI-derived features (T1w/T2w ratio, cortical thickness, sulcal depth, surface area, gray matter volume, mean curvature, gaussian curvature). This resulted in a single MIND matrix per subject, with brain regions represented as columns and rows. From each subject’s MIND matrix, we computed a regional connectivity strength vector (i.e., total similarity strength) by taking the mean across each row of the MIND matrix. This vector represents the average morphometric similarity of one region to all other regions for a given subject. Two metrics of interest were calculated. (1) The cohort-level total similarity strength was constructed by averaging the subject-level total similarity strength vectors and captures inter-regional relationships across all subjects, minimizing influence of age, while also amplifying signal-to-noise ratio. (2) Mixed effects models were fit for each region to assess age-related effects, with sex and hemisphere included as random effects. Age effects were quantified as the t-values from each regional model. Ex Vivo Transcriptomics. (bottom left) Cell type specific distributions mapped on the left hemisphere of the macaque cortex. Figure adapted from Chen et al. under a Creative Commons CC-BY license ^38^. (bottom right) The spatial distributions of 264 cell types were grouped into 23 broader categories. These were used to determine the underlying cell types that drive total similarity strength and age effects using CCA and Pearson correlation.

First, total similarity strength had significant negative correlations with glutamatergic cell types in layers 4/5 (L4/5; r = -0.27, p_spin_ = 0.017) and 5/6 (L5/6; r = -0.24, p_spin_ = 0.026) (Fig 2A). Conversely, oligodendrocytes (OLG), a glial cell-type that myelinates the axons of neighboring neuronal cells, had a positive correlation with the total similarity strength, but did not survive conservative correction (r=0.19, p=0.08). Oligodendrocytes are located primarily in the deep layers of cortex^38^, situated alongside the deep layer excitatory neurons.

**Figure 2:**
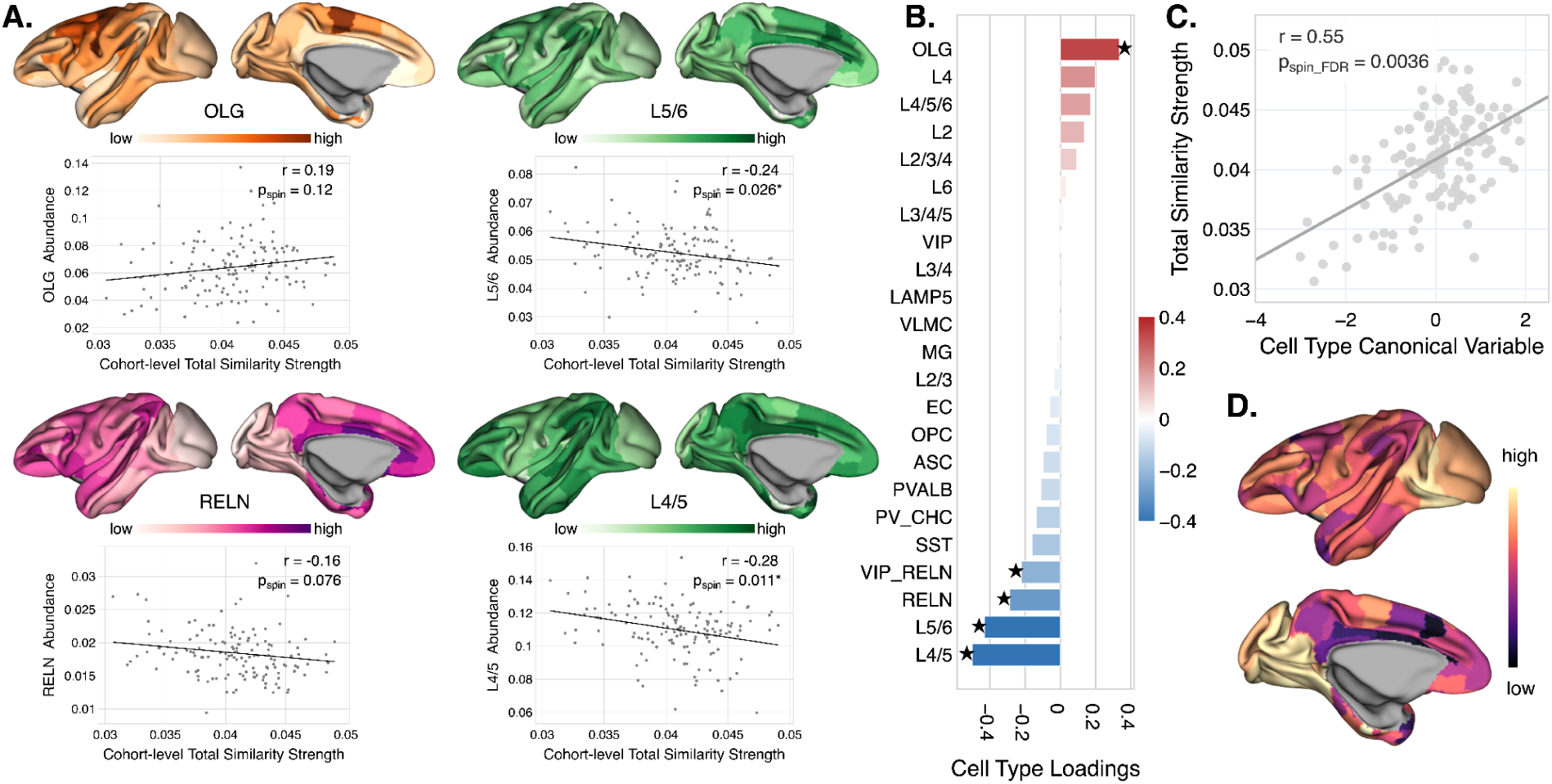
A. Scatterplots depict cell types with the strongest association with total similarity strength, with corresponding cell type abundances visualized on the left hemisphere CIVET macaque surface. Each Pearson correlation was evaluated with spin testing using 5,000 Hungarian null maps. B. Canonical Correlation Analysis (CCA) was performed between total similarity strength and cell type spatial distributions (r = 0.55, p_spin_FDR_ = 0.0036). Significance was assessed using 5,000 Hungarian null models. Cell-type loadings were defined as correlations with the cell type canonical variable. Significance was determined via two-sided spin testing with FDR correction. C. Total similarity strength plotted against the cell type canonical variable. D. Spatial distribution of the canonical cell variable visualized on the CIVET surface mesh.

Second, Canonical Correlation Analysis (CCA) was used to find a unique linear combination of cell type-specific abundances, here referenced as cell type canonical variable, that is maximally correlated with the total similarity strength, in a similar manner to ref ^3,44^ (Figure 2B-D). The magnitudes of relative contributions, or *loadings*, quantify the spatial association between the corresponding cell type abundances and canonical variable, while their signs indicate the directionality of such associations. Within the CCA model (r= 0.55, p_spin_FDR_ = 0.0036), we observed positive loadings of L4/5, L5/6, RELN (reelin), VIP (vasoactive intestinal peptide), VIP_RELN, and OLG . The findings were robust to stability testing unless OLG was omitted, thereby highlighting its contribution to the association.

### 2.2 Age-related differences in cortical similarity vary across functional networks in macaques

There is evidence from people and rats that multivariate cortical similarity networks are sensitive to age ^24,31,45^ , but whether that relationship holds in rhesus macaques has not been tested. Most prior studies typically focus only on early development or old age, making our lifespan coverage particularly unique.

We evaluated how age influences total similarity strength across the brain, within functional networks, and in individual regions in a cohort of healthy rhesus macaques (1 - 26 years). We grouped brain regions using a seven-network parcellation ^5^, reflecting standard definitions of fMRI-derived human functional networks that have evidence of conservation in macaque ^46–52^. For example, lower-order or unimodal areas are supported by the visual and somatomotor networks, where higher-order or transmodal networks, which mediate abstract thinking, memory and emotion, include the default mode and limbic networks ^4^. Such gradient hierarchy from unimodal to transmodal regions has been observed in humans and macaques, indicating this particular axis of functional and structural variation is conserved across species ^4,53^.

Mixed-effects models, on a whole brain, network and regional scale, were used to assess linear associations between age and total similarity strength, accounting for sex, subject, and cortical hemisphere, where appropriate. We observed a significant decrease in total similarity strength with age within the default mode, limbic, ventral attention, and frontoparietal networks and in the whole brain model, but not in the visual, dorsal attention, and somatomotor networks (Fig 3A-B). Notably, networks that were significantly impacted by aging aligned with higher order functional systems, while those without significant effects were associated with lower order sensory processing.

**Figure 3:**
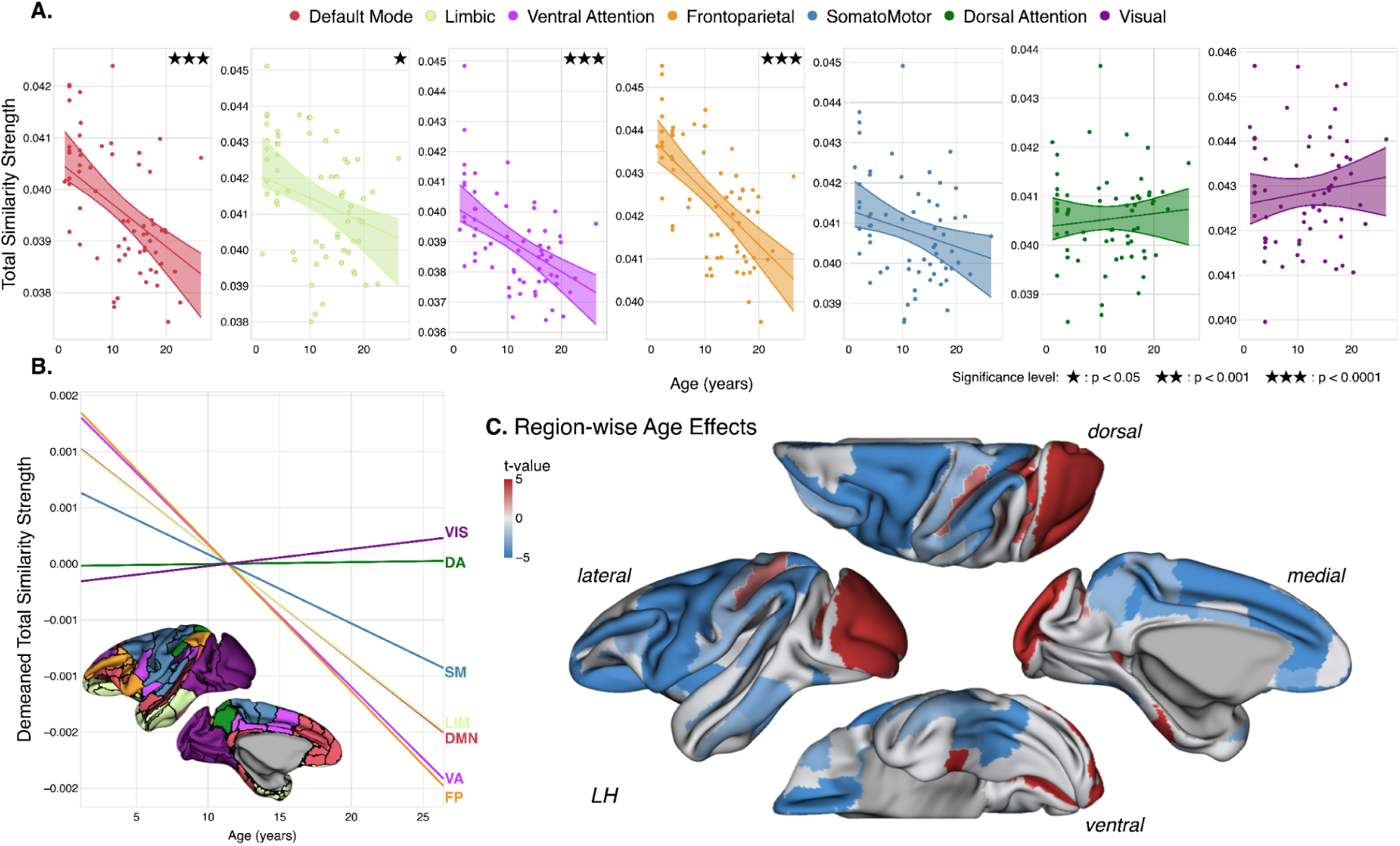
A. Age effects of total similarity strength are shown for large-scale functional networks as adapted from the Yeo parcellation ^5^ for macaque brain. Asterisks denote significant effects and shaded bars overlaid on each plot represent the 95% confidence interval of the estimated mean. Total similarity strength is defined as the mean similarity per region. B. To visualize age-related trends across networks, data were demeaned within each functional network and the slopes of the demeaned models were plotted. C. Regional age effects were evaluated using mixed effects models. Regions with significant age-related t-values (p_FDR_ < 0.05) are shown on the left hemisphere. Nonsignificant regions appear in light gray.

To assess potential heterogeneity at the level of smaller subregions within functional networks, we assessed the magnitude and direction of age-related effects across aggregate regions (Fig 3C, Supplementary Fig 1). Regions within sensory networks (visual, dorsal attention, and somatomotor) showed significant increases in total similarity strength with age. Within the dorsal attention network, these included MIP, LIPv, VIP, and PIP—all subregions of the intraparietal sulcus, an area associated with sensorimotor function. V3A, V3d, V1, and V6 in the visual network and 5_(PE) and 5_(PEa) in the somatomotor network, also exhibited positive relationships with age. Together, these regions are often implicated in the dorsal visual stream, a pathway involved in motion processing and spatial awareness^54^. Although the overall trend across each of the visual, dorsal attention, and somatomotor networks appears insignificant (Fig 3A), examining individual regions reveals a mix of positive and negative relationships with age, helping to explain the lack of a significant global effect.

### 2.3 Inhibitory and glial cell-types are associated with a decline in cortical similarity across the lifespan

The aging primate brain undergoes nonuniform morphological changes, which have been extensively characterized using structural MRI ^55–57^. The cellular mechanisms behind these changes remain understudied, especially in humans where acquiring brain tissue is much less feasible. Here, using cell type spatial distributions in the macaque cortex^38^, we investigated the cell type signatures driving age-related differences in cortical architecture to serve as a translational bridge to inform human aging research.

Region-wise age effects from the linear mixed effects models were correlated with the regional spatial distributions of each cell type. PV_CHC (r = -0.39, p_spin_ =0.006) was the only cell type that had significant negative associations with age effects (Fig 4A). RELN, which showed the second strongest negative association with age effects (r =-0.23, p=0.09), did not survive significance testing at conventional levels of significance. These inhibitory cell types are typically localized in more superficial layers of cortex, with PV_CHC expressed in layers 2 and 4, and RELN in layers 1 and 2 ^38^ . This result indicates that PV_CHC tends to reside in brain regions that exhibit decline in similarity strength due to age.

**Figure 4:**
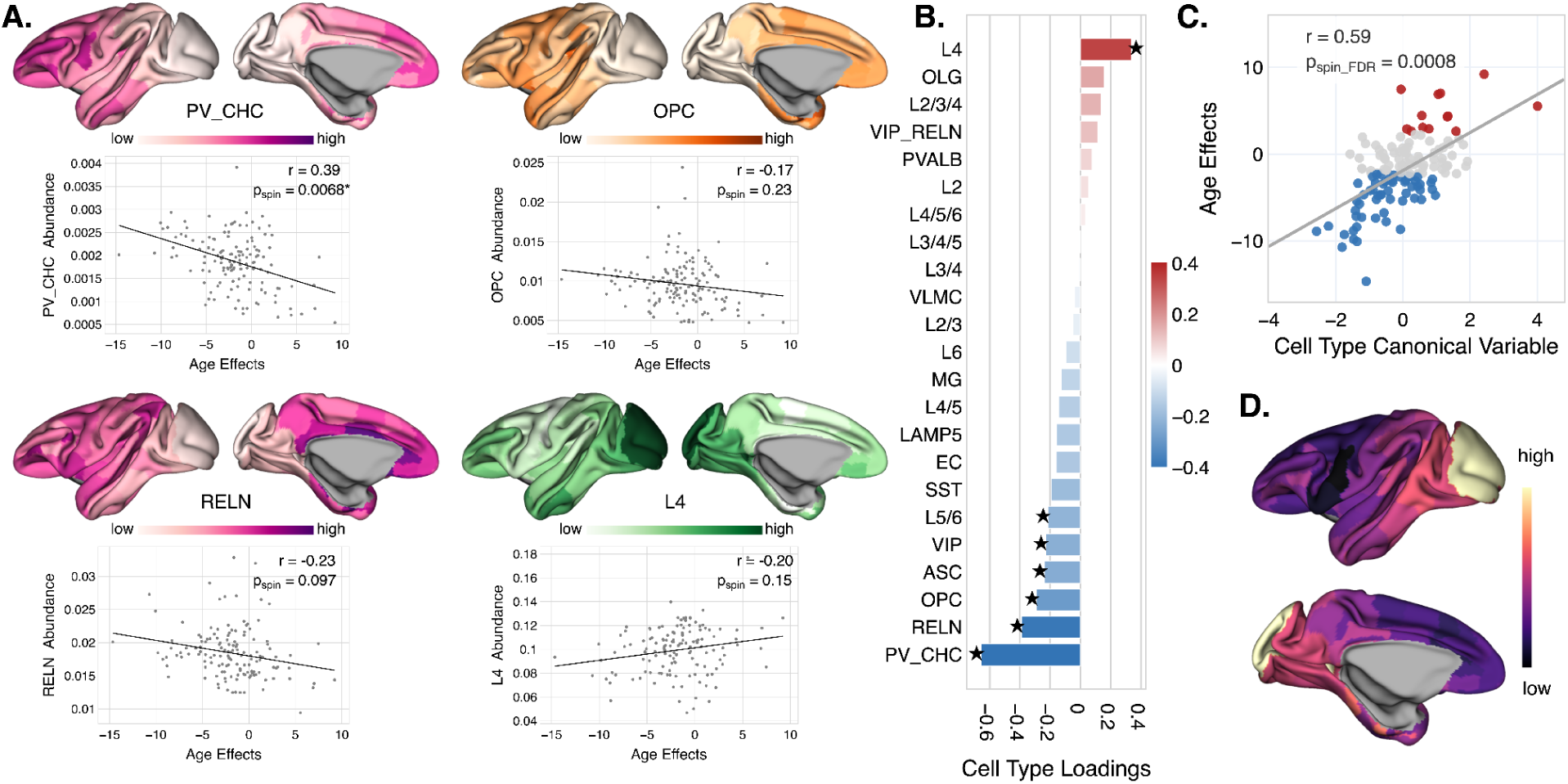
A.Scatterplots depict cell types with the strongest association with age effects, with corresponding cell type abundances visualized on the left hemisphere CIVET macaque surface. Each Pearson correlation was evaluated with spin testing using 5,000 Hungarian null maps. B. Canonical Correlation Analysis (CCA) was performed between age effects and cell type spatial distributions (r = 0.59, p_spin_FDR_ = 0.0008). Significance was assessed using 5,000 Hungarian null models. Cell-type loadings were defined as correlations with the canonical cell variable. Significance was determined via two-sided spin testing with FDR correction. C. Age effects plotted against the cell type canonical variable, with each point representing a unique region. Points are colored if they had significant age effects (see Fig 3C), with red showing significantly increased total similarity strength and blue showing significantly decreased total similarity strength with age. D. Spatial distribution of the cell type canonical variable visualized on the CIVET surface mesh.

When investigating the relationship of multiple cell types using CCA (r = 0.59, p_spin_FDR_ = 0.0008), age effects had significant loadings for PV_CHC, RELN, OPC, VIP, L5/6, ASC, and L4 (Fig 4B-D). Even when combinations of cell types with the most significant loadings were removed from the dataset in a stability test and a CCA was rerun, the association was still statistically significant. This indicates that the multivariate relationships were not being driven by a single cell type but rather the relationship between cell types. CCA loadings give the correlation of a cell’s spatial distribution to the cell type canonical variable, where a positive loading indicates that the cell is increasing in regions where the cell type multivariate profile is also increasing and a negative loading indicates the cell has an inverse pattern in comparison to this profile.

Both statistical approaches identified inhibitory cell types, including PV_CHC, RELN, and VIP, to be prominently expressed in brain regions that show significant decreases in similarity strength with age (Fig 4A). Notably, PV_CHC and RELN had the largest loadings from the CCA composite index. These cell types have higher relative enrichment in prefrontal and cingulate cortices, regions which are distributed across frontoparietal, default mode, and limbic network.

Other significant cell types, including non-neuronal cell types, OPC (oligodendrocyte progenitor cells) and ASC (astrocytes), and glutamatergic cell type L5/6, also had negative CCA loadings. The only significant positive loading came from the glutamatergic cell type L4.

### 2.4 Macaque cell type enrichment profiles diverge across higher and lower order cortical networks

Recent work in the human brain has shown that network features derived from resting state functional MRI exhibit preferential enrichment for particular neuronal and non-neuronal subtypes^3^. We extended this investigation to the nonhuman primate, using high-resolution spatial transcriptomic data from the macaque cortex. By quantifying cell type enrichment across canonical functional networks and applying hierarchical clustering, we aimed to identify network-specific profiles and their relationship to cortical hierarchy. An enrichment score (z-score) derived using spatial permutation testing, reflects how much a specific cell type is over- or underrepresented within each functional network, relative to its cortical distribution (Fig 5A).

**Figure 5:**
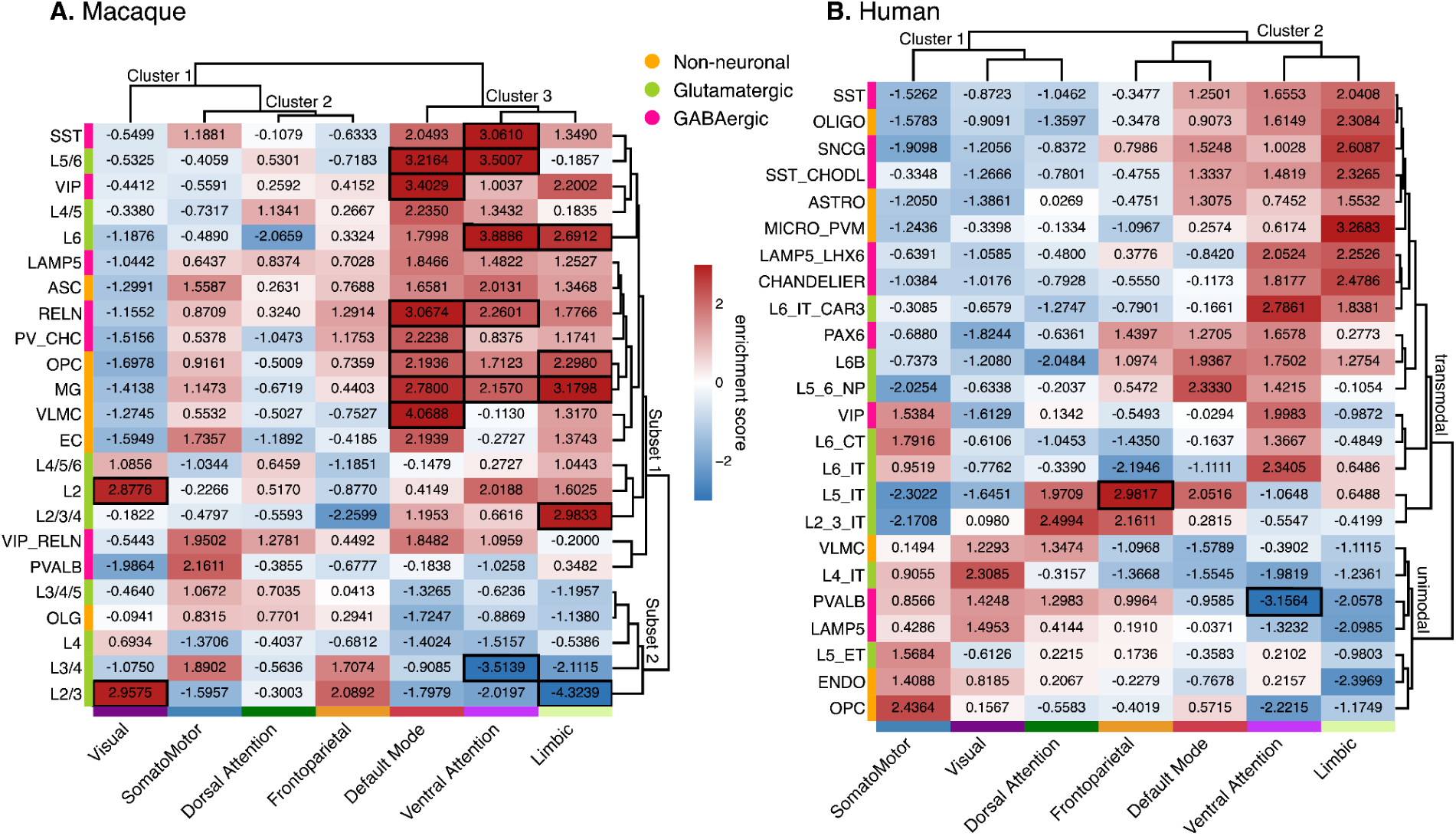
A. Enrichment scores (z-scores) of each cell type were computed across functional networks in the macaque cortex. D99 parcel values were projected to the surface and subjected to vertex-wise spin tests, generating 5,000 null maps per cell type. For each null map, vertex-wise cell densities were averaged within functional networks to create null distributions. Empirical enrichment scores were converted to z-scores using the null mean and standard deviation. P-values were computed as the proportion of null values more extreme than the empirical score and FDR-corrected across 23 cell types within each network. Significant enrichments (p_spin_FDR_<0.05) are outlined. B. Human functional network enrichment calculated from the estimated cell abundance from Zhang et al. ^3^ with 5,000 null maps per cell type. For each cell type, parcel-wise data in the Schaefer 400 atlas was projected onto vertices of a fsaverage6 sphere and random rotations were applied. Permuted vertex-wise data was grouped by functional network label and averaged to create null functional network maps. P-values were calculated by finding the proportion of null enrichment scores that were more extreme (two-sided) than the observed enrichment score per network. FDR correction was applied within each network separately. Enrichment scores annotate each entry and significant enrichment scores (p_spin_FDR_<0.05) are outlined in black.

Hierarchical clustering was performed to organize patterns within the enrichment matrix. Across networks (columns), clusters highlighted the divide between higher and lower order processing functions in the brain. The visual network is dominated by regions that receive sensory information, ranking it low in the functional hierarchy of visual processing ^58^. It did not align with any other functional domain (Cluster 1), and showed a remarkable inverse pattern of cell type enrichment compared to the three higher order networks. Within the visual network, which had relatively low enrichment of most cells, several cell types stood out with significant positive enrichment: L4/5/6, L2, L4, and L2/3, with L2 and L2/3. Interestingly, L4 and L2/3 neurons are often noted for their role in the visual processing pathway.^59–61^

Cluster 2 included the somatomotor, dorsal attention, and frontoparietal networks. Each of these networks had more heterogeneous patterns of expression, which may be why they were classified together. Most glutamatergic neurons, except for L3/4 and L3/4/5, exhibited negative enrichment in the somatomotor network. Among non-glutamatergic types, the VIP inhibitory cell type was the only cell that also had negative enrichment.

For higher order networks (Cluster 3), default mode, limbic, and ventral attention shared similar cellular enrichment profiles and they also contained the greatest number of significantly enriched cell types. Within the default mode network, there was a significant positive enrichment of L5/6, VIP, RELN, PV_CHC, OPC, MG (microglia), and VLMC (vascular leptomeningeal cells). Limbic had significant positive enrichment of L6, OPC, MG, and L2/3/4, and significant negative enrichment of L2/3. Ventral attention had significant positive enrichment of SST, L5/6, L6, RELN, and MG, and had significant negative enrichment of the L3/4 cell type.

Hierarchical clustering of cell types (rows) revealed two major subsets of cells based on a contrasting high / low pattern of enrichment across networks. The first and largest subset (Subset 1) showed exclusively positive enrichment in Network Cluster 3 (default mode, limbic, ventral attention) contrasted by mostly negative enrichment in Cluster 1 (visual). This grouping included all the GABAergic cell types, all non-neuronal cells except for OLG, and some glutamatergic cells in layers 2, 4-6. There was less clear alignment at the middle of the hierarchy (somatomotor, dorsal attention, frontoparietal).

Subset 2 cells had a less defined pattern of enrichment. It showed negative enrichment in Cluster 3 networks and consisted mainly of glutamatergic cell types from cortical layers 2–4. Some positive enrichment of Subset 2 appeared in visual and Cluster 2 networks.

### 2.5 Macaque and human cell type enrichment profiles have conserved network level patterns

To assess the cross-species generalizability of network-level cell type enrichment from the previous section, we compared spatial enrichment profiles derived from macaque transcriptomics data with those estimated in the human cortex after reanalyzing expression data from Zhang et al. ^3^ (Fig 5B, Methods Section 4.12). The human cortical dataset from Zhang et al. ^3^ contains cell type distributions imputed by applying cellular deconvolution to bulk gene expression data from the Allen Human Brain Atlas (AHBA)^62^, using cell type-specific signatures derived from single-nucleus RNA-seq data published by Jorstad et al. ^63^

Across species, the overall pattern of positive and negative enrichment was similarly organized along a shared cortical hierarchy: transmodal networks (e.g., default mode, limbic, ventral attention) showed enrichment of inhibitory and non-neuronal cell types, while unimodal networks (e.g., visual, somatomotor) were enriched for specific glutamatergic subtypes. This cross-species alignment reinforces the hypothesis that cellular specialization follows conserved principles of cortical organization, despite species-specific differences in data resolution, transcriptomic depth, and cell type labeling.

Hierarchical clustering of the enrichment matrices in both species revealed a separation of cell types, largely reflecting enrichment in higher order versus lower order processing (Fig 5). Zhang et al. identified these two subsets of cell type clusters (rows) as transmodal and unimodal ^3^. Our analysis revealed similar organizational structure, with some small differences in the group membership of cells exhibiting enrichment features of both cell clusters (Fig 5B). This is likely due to variations in clustering parameters.

Macaque Subset 1 had similar properties as the human transmodal cell cluster and included a mix of non-neuronal and inhibitory cell types. In contrast, macaque Subset 2 exhibited the opposite pattern, with negative enrichment in higher order networks and positive expression in lower order networks, resembling the human unimodal cell type cluster. We also qualitatively highlight similarities in cell type groupings across these parallel clusters, focusing on cell types that consistently co-occur (one to one) across species. Notably, several macaque cell types overlapped with the human transmodal cluster, including ASC, MG, SST, and PV_CHC. All of these cell types showed positive enrichment in higher-order networks in both species, suggesting a conserved direction of effect ^64,65^. The positive enrichment may reflect a higher baseline density of support and modulatory cells in the transmodal cortex, consistent with its later maturation and greater plasticity. No macaque cell types in Subset 2 directly matched the human unimodal cluster due to differences in taxonomy, gene panels, or sparse data in low-expression regions.

In the reanalyzed human dataset, enrichment profile trends by cell type and network were broadly consistent, demonstrating the stability of this method. Within our analysis, the frontoparietal network exhibited significant positive enrichment for L5/IT neurons, while the ventral attention network showed significant negative enrichment for PVALB-positive interneurons.

The macaque analysis showed more significant effects, but emphasis should be placed on effect direction and relative ordering, rather than the total amount of effects. This discrepancy in number of significant effects may be attributable to differences in atlas resolution, functional grouping assignments, or cell type annotation across species. Prior work has demonstrated that, within enrichment analyses, directionality is robust, even though significant partitions may vary ^66^.

### 2.6 Cross-species comparison of cell type composition aligns with the sensorimotor-association axis

To assess cross-species similarity in cortical cell type organization, we compared spatial enrichment profiles across functional networks in macaques and humans. Due to differences in how the gene expression data was acquired and ultimately labeled, a complete quantitative one-to-one cell type comparison was not possible. As a result, two analyses were performed to assess cross-species similarity. First, principal component analysis (PCA) was performed on each set of enrichment profiles separately to identify if dominant modes of variation in cell type composition align across species (for example, do cell types align with a principal organizational axis). Second, the Mantel test (see Methods Section 4.13) was performed to test if overall patterns of network similarity or dissimilarity exist between species (for example, how different is default mode from visual).

For PCA, there were three components that explained 58%, 19%, and 9% of the variance of the data in the macaque dataset and 56%, 24%, and 9% of the variance of the data in the human dataset. The first principal components from each PCA were found to be significantly correlated (r =0.85, p=0.01) with each other. Both of these latent variables were found to align with the hierarchy of functional networks in the cortex (Fig 6A). Further, when PC1 for each species was plotted onto their respective cortical surface, both aligned with the sensorimotor-association (S-A) axis (Fig 6A). This suggests that the primary mode of variation in cell type specific distributions in both macaques and humans follows the S-A axis, consistent with previous findings that show the alignment of various related neurobiological properties with this axis ^67–70^. The second (r=0.10, p=0.83) and third (r=0.67, p=0.10) principal components were not aligned between macaque and human cellular distributions.

**Figure 6.**
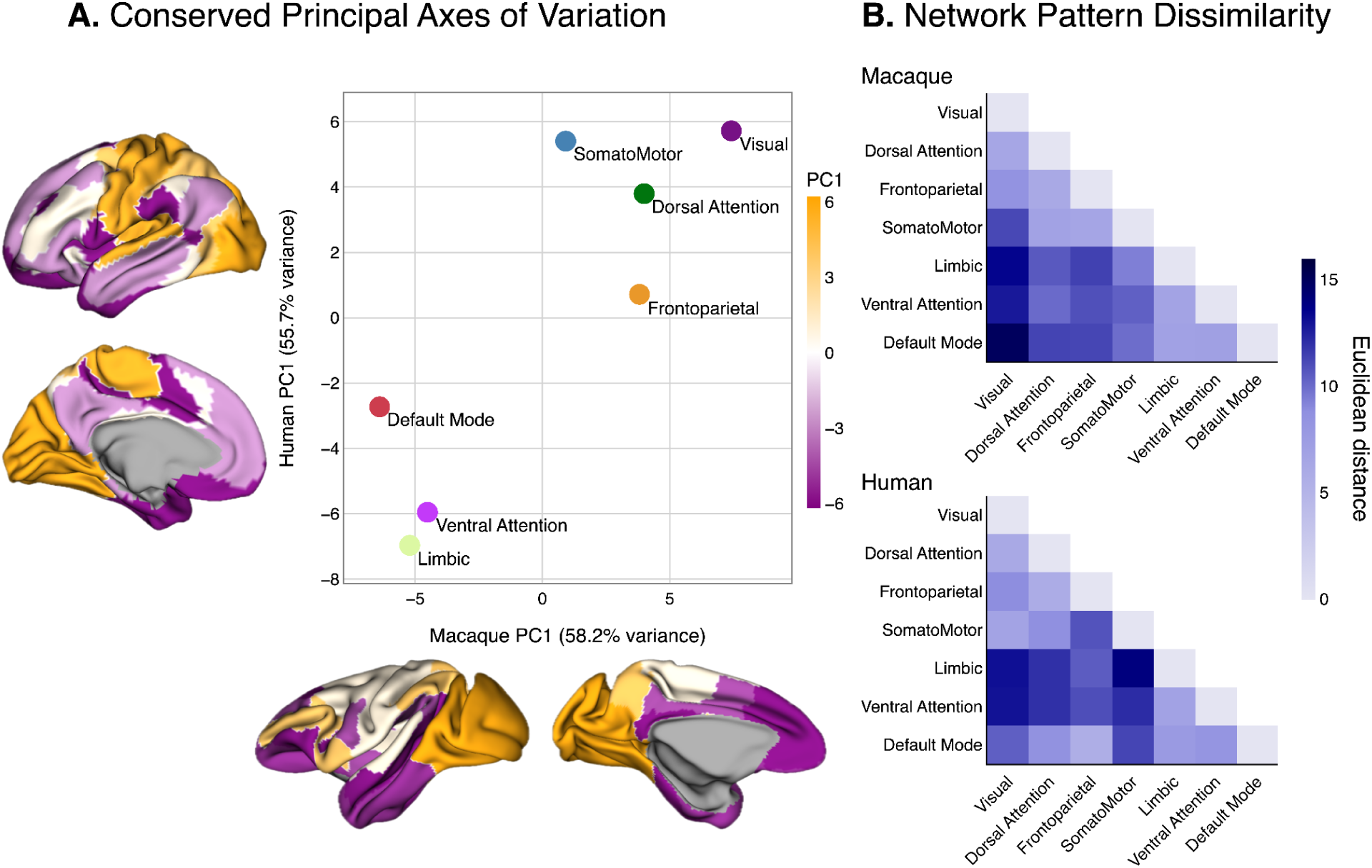
A. PCA was fit on each enrichment matrix separately. The first PC of the macaque dataset was plotted against the first PC of the human dataset, with percent of the variance explained given on the plot’s axes. Each of the first PCs were plotted on the human (top left) and macaque (bottom right) brain surfaces. The order of networks by PC1 in both primates aligns with the S-A axis. B. As a separate cross-species comparison, adjacency matrices based on Euclidean distances were generated to capture inter-network relationships in each primate. This approach allows for comparisons even when exact features (e.g., cell types) differ across datasets. The lower triangular (non-diagonal) values of each matrix were extracted and correlated to assess the similarity of network organization between humans and macaques (r = 0.47, p = 0.03).

Second, the Mantel test showed a significant cross-species correlation (r=0.48, p=0.027), suggesting that despite shifts in individual network pairings, the overall structure of cell type similarity is preserved. While specific close or distant relationships vary (Fig. 6B), broad trends between higher- and lower-order networks remain consistent.

The major differences in network relationships occurred in the position of the default mode and somatomotor networks within the hierarchy. Within the macaque, the default mode appeared to be the base of the functional hierarchy, as it contained the most over-enriched cell types and was ranked the most extreme by the first PC of cell types. In contrast as defined by the PCA, the human default mode was ranked third in the functional hierarchy, with the limbic network ranking first. In macaque, the somatomotor network was more closely aligned with the frontoparietal and dorsal attention networks, whereas in the human cell type spatial data, it was most similar to the visual network, as ordered by PC1.

This reorganization may reflect the evolutionary expansion of “higher order” cognitive functions in the human brain, whereby the default mode network shifted from a foundational role to a more integrated position supporting complex functionality like social cognition, self-referential processing, language, and abstract thought ^71,72^. Hierarchical rearrangement may reflect disproportionate expansion of association cortex in humans and tighter somatomotor-visual coupling could enable enhanced manipulative abilities as in our proficient tool use and fine motor capabilities ^53^.

## 3. Discussion

Here we present the first lifespan characterization of cortical similarity networks in the macaque brain, linking age-related changes to underlying cellular architecture using high-resolution spatial transcriptomics. By integrating in vivo MRI-based connectivity measures with postmortem transcriptomic data, we identify consistent associations between structural connectivity, age effects, and the spatial distribution of specific neuronal and non-neuronal cell types.

We found that deep-layer excitatory neurons (L4/5 and L5/6) and oligodendrocytes, both in infragranular cortical layers, are spatially aligned with a static, or average, representation of cortical similarity. While MRI lacks cellular specificity, these findings suggest that regional variation in MIND similarity reflects underlying microstructure. Multivariate analysis showed that oligodendrocytes, though not being independently significant, contribute dominantly in a multi-cellular context. This aligns with human transcriptomic data showing increased expression of oligodendrocyte and L5/6 neuron during development ^73,74^, reflecting protracted myelination ^75^. That the adult macaque profile emphasizes similar deep-layer types suggests these features are stable components of cortical architecture.

Given the spatial properties of MRI-based MIND, it is plausible that cells with prominent morphology, such as large somas, extensive dendritic arbors, or myelination, may disproportionately shape the similarity structure ^73^. These findings support the idea that morphometric similarity reflects not only cell type abundance but also the biophysical characteristics of specific cell classes.

Investigating structural similarity across the lifespan is valuable for understanding how cortical network organization evolves with age. The Structural Model, proposed by Helen Barbas and colleagues, predicts that similar cortical areas are more likely to be axonally connected than dissimilar areas ^76,77^. This theory has been demonstrated in macaques via tract tracing and in humans through coordinated neuroimaging analyses ^27–29^ . Neurotransmitter systems further support this theory, as their spatial distributions follow cortical similarity gradients.^9,67^.

In our cross-sectional lifespan macaque cohort, we observed an age-related decline in network connectivity, measured by total similarity strength. This reduction was most pronounced in higher-order networks - frontoparietal, ventral attention, default mode, and limbic. This suggests functional networks might become less specialized with age, consistent with human fMRI studies ^78^. Although longitudinal data are needed to confirm within-individual trajectories, our findings are consistent with the idea that higher-order networks are especially vulnerable to aging, reflecting a conserved pattern across primates.

This apparent disconnection may reflect microstructural and cellular changes that disrupt large-scale integration. Postmortem studies have investigated potential drivers, implicating synaptic integrity ^16^, laminar differences in cytoarchitecture, neuronal density^18^, and the density and targets of cortico-cortical projections. In *vivo* neuroimaging reveals cortical thinning, reduced surface area, and altered gyrification ^14,17^. Together, these findings suggest that morphometric similarity captures meaningful biological variation that may serve as a sensitive marker of aging-related vulnerability.

Inhibitory interneurons (PV_CHC, RELN, VIP) and non-neuronal cells (ASC, OPC) were enriched in cortical regions with the strongest age-related decline in cortical similarity, suggesting a potential link between metabolic demand and network vulnerability. This aligns with prior postmortem findings showing lower proportions of glial and inhibitory cells in older macaques ^19^. Among inhibitory subtypes, parvalbumin-positive chandelier cells (PV_CHC) showed the strongest spatial alignment with decline. These highly arborized, fast-spiking cells have high metabolic demands ^79,80^, and uniquely target the axon initial segment of pyramidal neurons, exerting strong inhibitory control ^81^. The high metabolic demands of these neurons, coupled with their late maturation ^81–83^ and known involvement in both neurodevelopmental and neurodegenerative disorders ^81,84–89^, suggests that parvalbumin expressing chandelier cells may play an important role in age-related changes of cortical networks. Although PV_CHCs are a relatively small subset of inhibitory neurons^38,90^, their distribution in macaques mirrors that in humans, suggesting conserved roles in cortical aging.^91^

Astrocytes and OPCs have distinct yet complementary roles in metabolic support, plasticity, and structural maintenance ^92–94^. These glial cells interact closely with inhibitory neurons, and aging-related astrocyte reactivity or OPC loss may disrupt this balance ^15^. Their shared alignment with vulnerable networks suggests that glia–interneuron interactions are a key axis of cortical aging.

Conversely, glutamatergic L4 neurons were the only cell type to have a positive age-related association. These neurons are most abundant in V1, reflecting specialized inputs coming from the thalamus for visual processing ^59–61,63^. V1 has a notably higher ratio of excitatory-to-inhibitory neurons, and was one of the few cortical regions that did not show age-related structural vulnerability in our data ^,63^. This suggests that L4 neurons may represent a preserved or selectively maintained population in this otherwise stable region.

Finally, we examined whether cell type enrichment profiles varied across functional networks, independent of MRI findings. We observed a conserved organization across species where both macaque and human cortex showed a principal axis of variation aligning with the sensorimotor–association gradient ^6,95^ . However, cross-species comparison was limited by differences in data type. Macaque Stereo-seq covered the whole cortex, while human data came from only 8 regions. Ideally, whole brain coverage of Stereo-seq or a related technology would be applied directly to human tissue. However, due to the significantly larger size of the human brain, applying Stereo-seq at scale remains costly and time-intensive. That said, efforts to process human brain tissue have already begun ^96,97^, opening the door for more precise cross-species comparisons in the future.

The spatial distributions of cell types can inform their role in aging and connectivity, particularly when they co-occur in vulnerable regions. However, aging and connectivity are complex, and spatial profiles alone do not capture cell morphology, physiology, or dynamic states. Moreover, condensing cell labels into 23 categories may obscure finer signals from subtypes with more nuanced spatial profiles. Despite these limitations, our findings suggest that cell type-specific distributions offer meaningful insights into cortical organization and age-related vulnerability.

In conclusion, these findings demonstrate a basis for aging-related changes in cortical similarity networks. Conserved cell type patterns across species support the macaque as a translational model for studying aging in humans. Advances in spatial transcriptomics will clarify links between cellular architecture and network vulnerability, informing mechanisms and potential interventions in brain aging.

## 4. Methods

### 4.1 Subjects

Sixty-four rhesus macaque monkeys with ages from 1 to 26 years old (age: mean ± SD = 11.37 ± 6.71, male: N=35) were included in this study. Macaques are estimated to age at roughly 3-4 times the rate that humans do^98^, reach sexual maturity on average between 3-4 years of age, and are considered geriatric after ∼18 years of age. As such, the monkeys in our sample span an age range equivalent to roughly 4 to 80+ years in humans. All monkeys were born and raised in small (housing 10 to 30 monkeys) or large (housing 80 to 120 monkeys) corrals at the California National Primate Research Center. Monkeys were socially housed at the time of MRI data acquisition, in pairs or small groups indoors, or large groups outdoors. Indoor housed monkeys were maintained on a 12 hour light/dark cycle with lights on at 6:00AM and lights off at 6:00PM. All monkeys were fed monkey chow twice daily, fresh fruit multiple times per week, and had *ad libitum* access to water. Prior to imaging data acquisition, monkeys received physical exams to ensure normative physiology. Some monkeys were subjects in previously published experiments^99–101,49,102^.

All imaging procedures were approved by the Institutional Animal Care and Use Committee at the University of California, Davis, which is accredited by the Association for Assessment and Accreditation of Laboratory Animal Care International (AAALAC). Animal care followed the guidelines outlined in the 2011 *Guide for the Care and Use of Laboratory Animals* by the Institute for Laboratory Animal Research.

### 4.2 Structural MRI

In vivo MRI data was acquired on a 3T Siemens Skyra scanner using a dedicated 8-channel radiofrequency monkey head coil at UC Davis. Animals were sedated by ketamine (5 mg/kg) and MRI compatible anesthesia machine was used to maintain sedation with 1.0-1.7% isoflurane. The animals were positioned in an MRI compatible stereotaxic frame in the sphinx position. The head coil was attached to stereotaxic using a custom coil holder.

From human studies, head motion is a known confound for MRI based analyses, where morphometric features, such as cortical volume and thickness, are systematically altered, leading to regionally specific biases ^103^. In this study, head movements were effectively eliminated by sedation and stereotaxic placement, enhancing sensitivity and reproducibility of the MRI analyses ^104,105^.

Structural images used in this study included (1) T1-weighted 3D Magnetization Prepared Rapid Gradient Echo (MP-RAGE) sequence (voxel-size: 0.3x0.3x0.6mm3; TE/TR=3.7/2500 ms; TI=1100 ms, FA= 7 degrees), and (2) T2-weighted sequence (non-interpolated voxel-size: 0.4x0.4x0.8mm3; TE/TR=308/3000ms; Echo train length =133ms; FA= 120 degrees).

### 4.3 Cortical surface-based analysis

In vivo structural T1w and T2w images were used for cortical surface extraction, as implemented in CIVET-Macaque ^106^. The cortical surface is represented by ∼41k vertices per hemisphere, allowing dense sampling across the brain for precise representation of cortical regions. A per-vertex correspondence between white matter and pial surfaces enables computation of morphometric cortical features. Individual subject surfaces were aligned to an average population template ^42,43^, allowing for vertex-based group and atlas level comparisons. The D99 macaque atlas ^107,108^ was used to group vertices into 141 regions per hemisphere (Fig 3B). One region on each hemisphere was removed due to having only one associated vertex, see Supplementary Table 1 for details on region assignment.

### 4.4 Cross-species alignment of functional domains

Seven canonical functional domains were defined across the macaque cortex. These domains were originally established using human resting state data ^5^, however, previous work has demonstrated cross-species alignment to macaques using joint-embedding gradients ^109^ (https://github.com/TingsterX/alignment_macaque-human). To extend this approach to our analysis framework, the cross-species Yeo surface (using fsLR) was registered and resampled to the CIVET surface space using Connectome Workbench ^110^. Most D99 atlas labels were nested within one of the seven functional network groupings. In regions where there was a D99 region mapping to two or more network groups, an expert anatomist (JC) reviewed and reassigned regions as needed. This assignment ensured that each D99 atlas region was associated with a single functional network identity, which was critical for subsequent analyses of cell type enrichment across different functional networks. The region assignments to functional networks are available in Supplementary Table 1.

### 4.5 Cortical similarity networks

MIND provides a robust measure of within-subject architectonic similarity between cortical areas based on multivariate distributions of vertex-wise MRI data for macro- and microstructural features between region pairs ^24^. For each subject, a MIND matrix was constructed from cortical features, including T1w/T2w ratio, cortical thickness, surface area, gray matter volume, sulcal depth, mean curvature and gaussian curvature. The similarity between nodal pairs was computed using the symmetrized Kullback-Leibler (KL) divergence, a measure of distributional similarity. To estimate the KL divergence, a k-nearest neighbors (k-NN) approach was employed, which allowed for the estimation of vertex-wise feature distributions per region, derived from MRI data. The symmetrized KL divergence was transformed to range from 0 to 1, with 0 representing low similarity and 1 representing high similarity. This transformation facilitates easier interpretation of the similarity values, where higher values indicate more similar nodes in terms of the MRI feature distributions. As a result, a 280x280 matrix is observed for each subject, where each entry corresponds to the similarity between a pair of brain regions. The source code used to create the MIND matrices was from the Github repository released with the original paper (https://github.com/isebenius/MIND).

### 4.6 Computation of network-based properties

For each subject, *total similarity strength* was calculated as the average value across the subject’s MIND matrix rows, excluding diagonal values. Each value in the resultant total similarity strength column vector (280 x 1) represents the average MIND similarity, or strength, between a node and all other nodes.

In addition, the subject-invariant regional strength properties were estimated by averaging all subject-wise total similarity strength values, aligned by region label. For later comparison with cellular spatial distributions which are measured on the left hemisphere only, corresponding regions on the left and right hemispheres were averaged to give one similarity value per region (140 x 1). The MIND values of the average left and right hemispheres were highly correlated (r = 0.98, p < 0.0001; Supplementary fig) and, in order to increase statistical power, the dataset was reduced as such.

### 4.7 Statistical modeling of age effects

To investigate age-related patterns of total similarity strength characteristics within the seven functional networks ^5^, linear mixed-effects modeling was applied using the R package lme4 ^111^, at a network and region scale (Fig 3). For the global and network-wise models, sex, subject ID and hemisphere side were added as random effects. For regional models, only the sex and hemisphere side were added as random effects. The false discovery rate (FDR) correction was applied to network and regional analyses. Age effects in subsequent analyses were defined as the region-wise t-values for total similarity strength mixed models.

### 4.8 3D single-cell transcriptomic atlas of macaque monkey

Cortical layer- and region-specific distributions of cell types were obtained from the first openly available single-cell transcriptomic atlas of the entire macaque monkey cortex (https://macaque.digital-brain.cn/spatial-omics) ^38^. This dataset includes transcriptomic profiles from approximately 1.5 million cells sequenced from the left hemispheres of two cynomolgus macaques. The molecular signatures of these classified cells were then used to identify cell types in Stereo-seq derived maps. The identified cell types were hierarchically categorized, with the highest level consisting of the three main brain cell types for glutamatergic, GABAergic cells, and non-neuronal cell types that were further subdivided into 23 specific subclasses (see Figure 1). Glutamatergic cell types included 10 layer-based (abbreviated L) subclasses: L2, L2/3, L2/3/4, L3/4, L3/4/5, L4, L4/5, L4/5/6, L5/6, and L6. The GABAergic cell types were labeled by five dominant marker genes: lysosome-associated membrane protein 5 (LAMP5), vasoactive intestinal peptide (VIP), reelin (RELN), parvalbumin (PV), somatostatin (SST). There were two additional subclasses that had distinct morphological or layer based properties while still having overlap in primary marker genes:VIP_RELN, and parvalbumin expressing chandelier cells (PV_CHC). The non-neuronal cell types were labeled to include marker genes for six subclasses: astrocytes (ASCs), oligodendrocyte precursor cells (OPCs), oligodendrocytes (OLGs), microglia (MG), endothelial cells (ECs), and vascular leptomeningeal cells (VLMCs).

These cell types are predominantly glial, with the exception of ECs and VLMCs, which are of vascular and mesenchymal origin, respectively. The spatial distributions of these cell types at each classification level were integrated into the D99 macaque histological atlas ^112^. We note that cynomolgus and rhesus macaques have highly similar brain structures and overall genetic similarity due to shared ancestry^113^ . Neuroscience laboratories regularly use both species interchangeably, and individual brains can be readily aligned between species. The D99 atlas, to which the single-cell transcriptomic data is registered, is indeed based on a rhesus macaque brain.

### 4.9 Univariate associations with cell type specific abundances

Using spatial permutation testing, the univariate associations were investigated between cell type specific abundance and our metrics of interest: the cohort level total similarity strength and age effects. For these analyses, the Hungarian method of creating spatially autocorrelated null maps was used ^66,114^. This algorithm generates null maps that are perfect permutations of the input data and was selected because it preserves the directionality of each value^66^, which was an important consideration for null maps of the age effects metric.

Parcel centroids from the D99 atlas were calculated using geodesic distance and projected onto the CIVET sphere. Random rotations were applied and the values of each parcel were optimally reassigned based using the Hungarian combinatorial optimization algorithm. 5,000 Hungarian null maps were created per metric of interest and used to create a distribution of null correlation values for each cell type. P-values were calculated as the proportion of null correlations that were more extreme (two-sided) than the true correlation.

### 4.10 Canonical Correlation Analysis

Canonical Correlation Analysis (CCA) is a technique used to understand and evaluate the multivariate cell spatial distribution relationship with regional metrics of interest. Analogous to a linear regression model, CCA determines linear combinations of two sets of variables that maximize their mutual correlation. CCA was conducted using PermCCA ^115^. Whereas the typical CCA is used to compare two groups of variables, in this study, the goal was to see how the multivariate cellular profiles interact with the metrics of interest (i.e., cohort-level total similarity strength, age effects) separately. As such, one CCA was performed per metric of interest.

Due to the spatial autocorrelation of brain surface data, for each regional metric of interest, 5,000 Hungarian null models were generated to evaluate the significance of the CCA.

Within each analysis, a canonical variable for cell density was produced, representing the linear combination of cell type-specific densities that exhibits the strongest correlation with the metric of interest. To determine the contribution of each cell type to the cell density canonical variable, loadings were computed by correlating each empirical cell density with the cell density canonical variable.

A sensitivity analysis was performed to determine whether certain cell types were dominating CCA results. Within this analysis, combinations of the cells with the most significant loadings per model were removed from the cell density dataset and a new CCA was computed. The significance of each resultant CCA was assessed using permutation testing with 5,000 Hungarian null maps.

### 4.11 Human specific cortical cell type enrichment dataset

The human cortical cell type imputed distributions from Zhang et al.^3^ were derived from snRNA-seq data across eight cortical areas in the Jorstad dataset^63^ and applied to AHBA bulk-tissue samples^62^ through deconvolution. For further details explaining the deconvolution methodology see ref ^3^.

The naming convention for the 24 human cell types follows that used in the Jorstad snRNA-seq dataset^63^. The nine GABAergic inhibitory interneurons, including PAX6 (paired box 6 gene), SNCG (synuclein gamma), VIP (vasoactive intestinal peptide), LAMP5 (Lysosomal-associated membrane protein family member 5 ), LAMP5_LHX6 (Lysosomal-associated membrane protein family member 5/LIM Homeobox 6), Chandelier, PVALB (parvalbumin), SST (somatostatin), and SST_CHODL (somatostatin/chondrolectin) are annotated by their marker genes.

The nine glutamatergic excitatory neurons, including L2/3 IT (intratelencephalic projecting), L4 IT, L5 IT, L6 IT, L5 ET (extratelencephalic projecting), L5/6 NP (near-projecting neurons), L6 CT (corticothalamic-projecting), and L6b are labeled using their layer preferences and their direction of projection. L6 IT Car3 (carbonic anhydrase 3) is also annotated by its marker gene.

Finally, the six non-neuronal cells are Astro (astrocytes), Endo (endothelial cells), VLMC (vascular and leptomeningeal cells), Oligo (oligodendrocytes), OPC (oligodendrocyte precursor cells) and Micro/PVM (microglia and perivascular macrophages).

### 4.12 Comparison of macaque and human cell types by their aligned functional networks

A cell type enrichment analysis was conducted to determine whether specific functional networks in the macaque cortex preferentially expressed certain cell type profiles. A second human cell type enrichment analysis was performed using the imputed spatial distributions of cortical cell type abundances, released by Zhang et al. ^3^. Both enrichment analyses followed the same methodology as follows.

The Cornblath method was used to generate 5,000 spatial null maps for each cell type ^116,117^. This method was selected because it provides a more conservative approach to null model generation and was recommended by Markello and Mišić ^66^. Furthermore, preserving the directionality of the null values was no longer a concern as it had been with previous null models.

The Cornblath null maps were created by projecting the empirical parcel-wise cell abundance data onto the vertices of a CIVET (macaque) or fsaverage6 (human) sphere, applying a random rotation, then projecting the data back onto the original surface and combining by functional network label. This process resulted in a null distribution of cell abundance estimates (5,000 null values) for each functional network label.

For each cell type, the network-specific mean and standard deviation were calculated. These values were then used to compute the enrichment score (z-score) of the original, unpermuted data and to standardize the network-wise distributions for p-value calculation. P-values were determined as the proportion of null enrichment scores that were more extreme (two-sided) than the true enrichment score. Hierarchical clustering was performed on the cellular profiles and functional profiles of the resultant cell type by functional network enrichment matrix.

To guide interpretation of enrichment analyses, prior work by Markello and Mišić comparing spatial null models shows that the number and identity of partitions with significant effects can vary across models, but the directionality of these effects are conserved across null model types and partitions^66^.

### 4.13 Macaque to human comparison of cell type composition

A principal component analysis (PCA) was performed on each enrichment matrix separately, treating functional network labels as observations. This was done to determine the axis of greatest variation, which we had hypothesized may align with the canonical functional hierarchy of both macaque and human cell type specific distributions.

Additionally, a Mantel Test ^118^ was conducted. This test assesses the similarity between two matrices with non-overlapping observations by comparing the structure of relationships among their columns. First, pairwise Euclidean distance matrices were computed to capture the dissimilarity between functional networks based on their cell type distributions. Their non-diagonal upper triangular values were aligned and correlated.

## 5. Contributions and Acknowledgements

### Author Contributions

E.P.R. and M.T. conceptualized the project and wrote the original draft of the manuscript. M.T. performed primary statistical analyses and visualizations. J.A.C. contributed anatomical expertise. J.L.B., J.A.C., E.B-M. acquired the in vivo MRI data. E.P.R, C.L., J.A.C., and J.V. preprocessed and analyzed the MRI data. J.V. provided supervision on imaging and statistical analyses. A.C.E. contributed funding and supervision. E.B-M. raised funding for all in vivo MRI data and provided supervision. E.P.R. contributed project funding and provided supervision. M.T., J.A.C., C.L., J.L.B., J.V., A.C.E., E.B-M., E.P.R. reviewed and edited the manuscript.

### Funding

This work was supported by the National Institute on Aging of the National Institutes of Health (NIH) under grant number R21AG083539 to EPR. JV is additionally supported by R21AG087904. In vivo MRI data acquired with support from K99MH101338, R01HD096436, and RF1AG078340, and the UC Davis Chancellor’s Fellowship awarded to E.B-M, and carried out at the California National Primate Research Center (P51OD011107).

## Acknowledgements

The results generated from CIVET-macaque draw on research supported by the Digital Research Alliance of Canada and the Government of Canada. We thank Xihan Zhang and Dr. Avram Holmes for valuable discussions towards developing the cross-species comparison.

## Data and Code Availability

Cell type specific spatial distributions of the macaque cortex were obtained from Chen et al.^38^ and can be found at https://macaque.digital-brain.cn/spatial-omics/singleCellData?project=neocortex. Imputed human cell type distributions were obtained from Zhang et al.^3^

All code generated for this study are available on github at https://github.com/micmaclab/celltype-primate-aging. Functional network alignment analyses were conducted using code from https://github.com/TingsterX/alignment_macaque-human. Spatial permutation tests utilized the netneurotools, nibabel, and neuromaps Python libraries. MIND network construction used code available at https://github.com/isebenius/MIND and canonical correlation analysis (CCA) was performed using https://github.com/andersonwinkler/PermCCA. Linear mixed effects models were implemented using the lme package in R. Surface visualizations were generated using https://github.com/danjgale/surfplot and Connectome Workbench.

