## Supplementary Information for "Cellular basis for cortical network aging in primates"

| Region | Name | Abbreviation | Label |
|---|---|---|---|
| 2 | dorso-lateral_prefrontal_area | 8Bm | Default Mode |
| 3 | orbital_prefrontal_area | 11l | Limbic |
| 5 | orbito-medial_prefrontal_area_(gyrus_rectus) | 14r | Limbic |
| 7 | agranular_insula | Ia | Ventral Attention |
| 8 | orbital_prefrontal_area | 13b | Limbic |
| 10 | area_36_of_the_perirhinal_cortex_caudal_subregion | 36c | Default Mode |
| 11 | subregion_of_anterior_cingulate_cortex | 24a' | Frontoparietal |
| 14 | medial_prefrontal_area | 10mc | Default Mode |
| 15 | caudal_parabelt_region_of_the_auditory_cortex | CPB | SomatoMotor |
| 17 | subregion_of_posterior_cingulate_cortex | v23b | Default Mode |
| 18 | orbital_prefrontal_area | 13l | Limbic |
| 20 | visual_area_4_(dorsal_part) | V4 | Visual |
| 23 | posterior_cingulate_area | 31 | Ventral Attention |
| 24 | granular_insula | Ig | SomatoMotor |
| 25 | middle_lateral_belt_region_of_the_auditory_cortex | ML | SomatoMotor |
| 27 | subregion_of_posterior_cingulate_cortex | 23a | Default Mode |
| 28 | entorhinal_cortex_rostral_division | ER | Limbic |
| 29 | orbital_prefrontal_area | 10o | Limbic |
| 30 | ventral_intraparietal_area | VIP | Dorsal Attention |
| 31 | lateral_intraparietal_area_ventral_subdivision | LIPv | Dorsal Attention |
| 32 | dorso-lateral_prefrontal_area_in_the_arcuate_sulcus_upper_limb | 8Bs | Frontoparietal |
| 34 | visual_area_1_(primary_visual_cortex) | V1 | Visual |
| 35 | gustatory_cortex | G | Ventral Attention |
| 37 | dorso-lateral_prefrontal_area | 9d | Ventral Attention |
| 38 | parietal_operculum | 7op | Ventral Attention |
| 39 | V4_transition_area | V4t | Visual |
| 41 | ventro-lateral_prefrontal_area_in_the_ventral_bank_of_the_prinicipal_sulcus | 46f | Frontoparietal |
| 43 | superior_parietal_lobule_area_5_(PE) | 5_(PE) | SomatoMotor |
| 44 | ventral_subregion_of_anterior_TE | TEav | Limbic |
| 45 | medial_prefrontal_area_32 | 32 | Default Mode |
| 46 | anterior_lateral_belt_region_of_the_auditory_cortex | AL | SomatoMotor |
| 47 | medial_prefrontal_area | 10mr | Default Mode |
| 49 | subregion_of_anterior_cingulate_cortex | 24b | Default Mode |
| 50 | somatosensory_areas_1-2 | 1/2 | SomatoMotor |
| 51 | periarcuate_area_(or_Frontal_Eye_Fields)_ventral_subdivision | 8Av | Frontoparietal |

| Region | Name | Abbreviation | Label |
|---|---|---|---|
| 52 | entrorhinal_cortex_caudal_limiting_division | ECL | Limbic |
| 53 | floor_of_superior_temporal_area | FST | Default Mode |
| 55 | orbital_prefrontal_area | 12m | Default Mode |
| 56 | visual_area_6A_ventral_subdivision_(or_PO) | V6Av | Frontoparietal |
| 57 | dorso-lateral_prefrontal_area | 8Bd | Default Mode |
| 59 | dysgranular_part_of_the_ventral_temporal_pole | TGvd | Limbic |
| 60 | visual_area_3_dorsal_part | V3d | Visual |
| 61 | agranular_frontal_area_F1_(or_4) | F1_(4) | SomatoMotor |
| 62 | medial_agranular_insula | Iam | Limbic |
| 63 | secondary_somatosensory_cortex | SII | Ventral Attention |
| 64 | subregion_of_posterior_cingulate_cortex | 23b | Ventral Attention |
| 65 | orbital_prefrontal_area | 13m | Limbic |
| 66 | dorso-lateral_prefrontal_area | 9m | Default Mode |
| 67 | entrorhinal_cortex_intermediate_division | EI | Limbic |
| 68 | rostral_parabelt_region_of_the_auditory_cortex | RPB | SomatoMotor |
| 69 | orbital_prefrontal_area | 13a | Limbic |
| 70* | anterior_olfactory_nucleus* | AONd/m* | Limbic* |
| 71 | agranular_frontal_area_F3_(or_SMA) | F3 | Ventral Attention |
| 72 | rostral_core_region_of_the_auditory_cortex | R | SomatoMotor |
| 73 | visual_area_6_(or_PO) | V6 | Visual |
| 74 | area_TFO_of_the_parahippocampal_cortex | TFO | Default Mode |
| 75 | visual_area_4_ventral_part | V4v | Visual |
| 76 | ventro-lateral_prefrontal_area_in_the_ventral_bank_of_the_prinicipal_sulcus | 46v | Frontoparietal |
| 77 | superior_parietal_lobule_area_5_(PEc) | 5_(PEc) | Dorsal Attention |
| 78 | dysgranular_part_of_the_dorsal_temporal_pole | TGdd | Limbic |
| 79 | area_35_of_the_perirhinal_cortex | 35 | Limbic |
| 80 | primary_auditory_cortex | A1 | SomatoMotor |
| 81 | lateral_agranular_insula | Ial | Limbic |
| 82 | 7_(PGm)_on_the_medial_wall | 7m_(PGm) | Dorsal Attention |
| 84 | visual_area_2 | v23b? | Default Mode |
| 85 | subregion_of_posterior_cingulate_cortex | 23c | SomatoMotor |
| 86 | entrorhinal_cortex_caudal_division | EC | Limbic |
| 87 | dysgranular_insula | Id | Ventral Attention |
| 89 | medial_intraparietal_area | MIP | Dorsal Attention |
| 90 | posterior_intraparietal_area | PIP | Dorsal Attention |
| 91 | rostral_inferior_parietal_lobule_area | 7b_(PFG/PF) | Dorsal Attention |

| Region | Name | Abbreviation | Label |
|---|---|---|---|
| 92 | dorsal_subregion_of_posterior_TE | TEpd | Limbic |
| 93 | visual_area_3_ventral_part | V3v | Visual |
| 94 | ventro-lateral_prefrontal_area_in_the_arcuate_sulcus_lower_limb | 44 | Default Mode |
| 95 | visual_area_MT | MT | Dorsal Attention |
| 96 | sts_ventral_bank_area | TEm | Limbic |
| 97 | temporo-parietal_area | Tpt | Ventral Attention |
| 98 | subregion_of_anterior_cingulate_cortex | 24c | Default Mode |
| 99 | visual_area_MST | MST | Default Mode |
| 100 | rostromedial_belt_region_of_the_auditory_cortex | RM | SomatoMotor |
| 101 | rostral_superior_temporal_gyrus | STGr | Default Mode |
| 102 | parainsular_area | Pi | Limbic |
| 103 | agranular_frontal_area_F6_(or_preSMA) | F6 | Default Mode |
| 107 | ventro-lateral_prefrontal_area | 12r | Default Mode |
| 108 | precentral_operular_area | PrCO | Default Mode |
| 109 | superior_parietal_lobule_area_(cingulate_sulcus) | PEci | SomatoMotor |
| 110 | granular_part_of_the_dorsal_temporal_pole | TGdg | Limbic |
| 111 | anterior_intraparietal_area | AIP | Dorsal Attention |
| 112 | area_36_of_the_perirhinal_cortex_temporal-polar_subregion | 36p | Limbic |
| 113 | lateral_occipital_parietal_area | LOP | Dorsal Attention |
| 114 | orbital_prefrontal_area | 11m | Limbic |
| 115 | polar_rostrotemporal_cortex | RTp | Limbic |
| 116 | anterior_olfactory_nucleus | AONl | Limbic |
| 117 | medial_prefrontal_area_25_(subgenual_cortex) | 25 | Limbic |
| 120 | rostrotemporal_core_region_of_the_auditory_cortex | RT | Limbic |
| 121 | caudal_inferior_parietal_lobule_area | 7a_(Opt/PG) | Frontoparietal |
| 122 | ventral_subregion_of_posterior_TE | TEpv | Default Mode |
| 123 | visual_area_V3A | V3A | Visual |
| 124 | ventro-lateral_prefrontal_area | 45a | Default Mode |
| 125 | area_TEO | TEO | Visual |
| 126 | agranular_frontal_area_F7_(or_6DR) | F7 | Default Mode |
| 127 | dorso-lateral_prefrontal_area_in_the_dorsal_bank_of_the_prinicipal_sulcus | 46d | Frontoparietal |
| 128 | intermediate_agranular_insula | Iai | Limbic |
| 129 | agranular_frontal_area_F5_(or_6Va/6Vb) | F5_(6Va/6Vb) | Frontoparietal |
| 130 | lateral_intraparietal_area_dorsal_subdivision | LIPd | Dorsal Attention |
| 131 | visual_area_2 | V2 | Visual |

| Region | Name | Abbreviation | Label |
|---|---|---|---|
| 132 | area_36_of_the_perirhinal_cortex_rostral_subregion | 36r | Limbic |
| 133 | ventro-lateral_prefrontal_area | 12l | Default Mode |
| 134 | superior_parietal_lobule_area_5_(PEa) | 5_(PEa) | SomatoMotor |
| 135 | sts_part_of_the_temporal_pole | TGsts | Limbic |
| 136 | ccg | RTM | Limbic |
| 137 | somatosensory_areas_3a_and_3b | 3a/b | SomatoMotor |
| 140 | agranular_part_of_the_temporal_pole | TGa | Limbic |
| 141 | visual_area_6A_dorsal_subdivision | V6Ad | Dorsal Attention |
| 142 | area_TF_of_the_parahippocampal_cortex | TF | Default Mode |
| 143 | ventro-lateral_prefrontal_area_in_the_arcuate_sulcus_lower_limb | 45b | Frontoparietal |
| 144 | dorsal_subregion_of_anterior_TE | TEad | Limbic |
| 145 | sts_fundus_area | PGa | Default Mode |
| 146 | ventral_premotor_cortex_(F4) | F4 | Ventral Attention |
| 148 | periarcuate_area_(or_Frontal_Eye_Fields)_dorsal_subdivision | 8Ad | Frontoparietal |
| 149 | lateral_rostrotemporal_belt_region_of_the_auditory_cortex | RTL | SomatoMotor |
| 150 | orbital_prefrontal_area | 12o | Default Mode |
| 151 | caudomedial_belt_region_of_the_auditory_cortex | CM | SomatoMotor |
| 152 | sts_dorsal_bank_area | TAa | Default Mode |
| 153 | agranular_frontal_area_F2_(or_6DR/6DC) | F2_(6DR/6DC) | SomatoMotor |
| 154 | granular_part_of_the_ventral_temporal_pole | TGvg | Limbic |
| 155 | caudal_lateral_belt_region_of_the_auditory_cortex | CL | SomatoMotor |
| 159 | sts_dorsal_bank_area | TPO | Default Mode |
| 160 | sts_ventral_bank_area | TEa | Limbic |
| 165 | sts_fundus_area | IPa | Limbic |
| 166 | entrorhinal_cortex_lateral_division_(rostral_part) | ELr | Limbic |
| 172 | visual_area_2 | V2? | Visual |
| 174 | visual_area_2 | V2_or_v23b? | Visual |
| 184 | area_30_(retrosplenial_cortex) | 30 | Default Mode |
| 185 | subregion_of_anterior_cingulate_cortex | 24b' | Ventral Attention |
| 187 | subregion_of_anterior_cingulate_cortex | 24c' | Ventral Attention |
| 188 | orbito-medial_prefrontal_area_(gyrus_rectus) | 14c | Limbic |
| 194 | entrorhinal_cortex_olfactory_division | EO | Limbic |
| 195 | area_TH_of_the_parahippocampal_cortex | TH | Limbic |
| 218 | posterior_lateral_agranular_insula | Iapl | Frontoparietal |
| 224 | retroinsula | Ri | SomatoMotor |

Supplementary Table 1. Assignment of D99 Regions to the Yeo 7-Network Parcellation. The Yeo 7-Network parcellation for macaque[110], originally in fslr space, was registered to the CIVET macaque surface. To align it with the D99 parcellation, the proportion of vertices within each D99 region corresponding to each Yeo network label was calculated. For regions where vertex proportions were similar across multiple Yeo labels, final assignments were made by expert anatomist JC. The Anterior Olfactory Nucleus (region 70) was removed in any further analyses due to having only one associated vertex in the CIVET macaque surface. Indicated with * in the table.

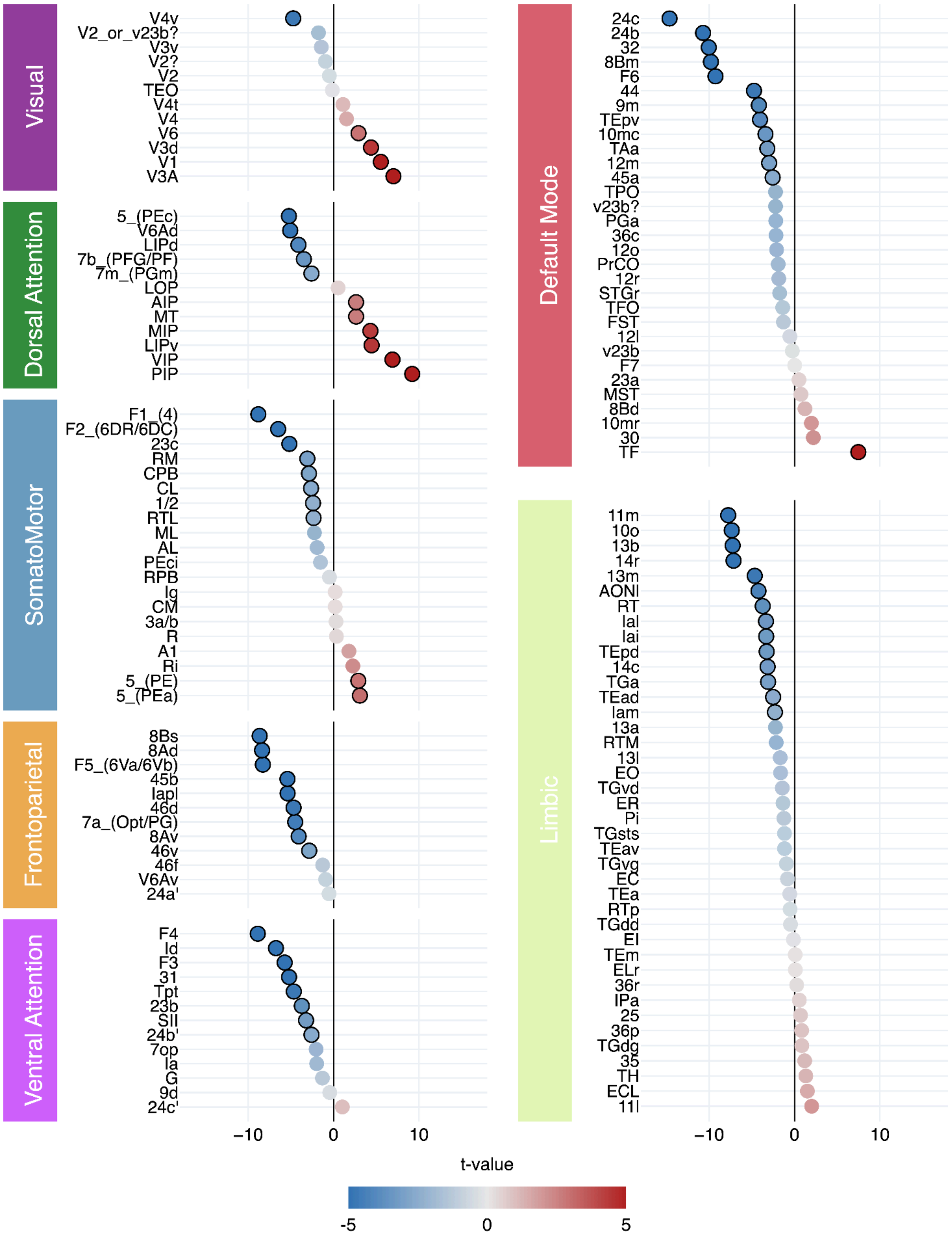


Supplementary Fig 1. Total similarity strength was used to create a linear mixed model for each region of the macaque brain. Points circled in black are significant ($p<0.05$) after FDR correction. Most network-wise trends are conserved at the regional level, except for several regions in the visual, dorsal attention, and somatomotor networks, which have a positive relationship with age.
